# The C-terminal regions of TRAK proteins contain MIRO-independent mitochondrial outer membrane binding domains

**DOI:** 10.1101/2021.06.03.446977

**Authors:** Lili Mitchell, Kathryn E. Reda, Hijab Fatima, Claudia E. Vasquez, Omar A. Quintero-Carmona

## Abstract

Current models suggest that MIRO GTPases anchor cytoskeletal motors to the mitochondrial outer membrane (MOM). However, our previous findings indicate that the unconventional myosin, MYO19, interacts with MIRO weakly and that a MIRO-independent MOM-localizing domain interacts more tightly with the MOM. To test the hypothesis that other MIRO interactors may also have MIRO-independent MOM-binding, we examined interactions between TRAK proteins (microtubule motor-mitochondria adaptor proteins) and the MOM via quantitative fluorescence microscopy and steady-state kinetic approaches. Using GFP-TRAK truncations expressed in MIRO1-2 double knockout mouse embryonic fibroblasts, we identified a MIRO-independent mitochondrial binding domain in the C-terminus of TRAK1 and TRAK2, sufficient for MOM-localization similar to what we observed for full length GFP-TRAK proteins. The MIRO-binding domains (MBD) of the TRAK proteins were only able to localize to mitochondria in the presence of ectopic expression of MIRO. Importantly, fluorescence recovery after photobleaching (FRAP) demonstrated that the steady-state kinetics of TRAK^MBD^/MIRO interactions were faster-exchanging than for either full-length TRAK or the TRAK C-terminal MOM-binding domain expressed alone. These data support a model where faster-exchanging TRAK/MIRO associations could support initial association and/or TRAK activation, while MIRO-independent binding contributes significantly to tighter association to the MOM.

## Introduction

Mitochondria serve many biological roles. In addition to adenosine triphosphate (ATP) synthesis, mitochondria participate in intracellular calcium homeostasis, regulate reactive oxygen species, and are central to apoptotic activities (Wang and Youle, 2009). The proper positioning of mitochondria within cells is essential for these activities and for maintaining cellular health. Improper localization can contribute to abnormal mitochondrial functioning which may contribute to degenerative diseases, metabolic diseases, and cancer (Wallace, 1999). Mitochondria positioning defects are implicated in neuronal dysfunction linked to Alzheimer’s and Parkinson’s disease (DiMauro, 1999). The study of mitochondrial localization, network organization, and transport are therefore important in providing foundational understanding of biological function and well as for developing treatment options for many serious diseases.

Mitochondria are dynamic organelles which are constantly dividing, fusing, and moving throughout the cytosol (Fenton et al., 2020). Mitochondria localize to different regions of the cell through interactions with motor proteins like kinesin, dynein, and MYO19. Attachment of these motors to mitochondria enable movement across cytoskeletal networks of microtubules and actin filaments. Actin-based mitochondrial movement is facilitated by interactions with a mitochondria-associated unconventional myosin, MYO19 (Quintero et al., 2009), while microtubule based motility is regulated by kinesins (Smith et al., 2006) and dynein (Pilling et al., 2006; van Spronsen et al., 2013). Many recent reports have implicated the small Ras-like GTPase, MIRO, as a receptor for motor proteins on the mitochondrial outer membrane (Glater et al., 2006; Guo et al., 2005; MacAskill et al., 2009; van Spronsen et al., 2013; Wang and Schwarz, 2009). Two isoforms of MIRO exist in mammalian cells, MIRO1 and MIRO2 (also known as RHOT1 and RHOT2). Human MIRO proteins consist of approximately 618aa and share ∼60% sequence identity. Both isoforms consist of an N-terminal Ras-like GTPase domain followed by two calcium-binding EF hand-like domains, a C-terminal Ras-like GTPase domain, and a transmembrane insertion domain. In addition to nucleotide state, MIRO activity is thought to be influenced by phosphorylation, and calcium (reviewed in (Eberhardt et al., 2020). A number of different binding partners have been shown to interact with MIRO through a hydrophobic region in in the first EF hand-like region (Covill-Cooke et al., 2024).

Microtubule motor protein interactions with MIRO on the MOM involve adaptors. Originally identified in *Drosophila*, TRAK proteins (also known as Milton) were thought to facilitate binding between MIRO proteins and microtubule-based motors (Stowers et al., 2002). Two mammalian TRAK proteins have been identified, TRAK1 (aka OIP106) and TRAK2 (aka GRIF-1), which share ∼58% amino acid homology (Beck et al., 2002; Iyer et al., 2003). The study of MIRO in a mitochondrial context has historically centered around its role in anchoring TRAK and molecular motors to the MOM (Fransson et al., 2003; Fransson et al., 2006). Though the precise mechanisms of MIRO, TRAK, and motor protein interactions have not been completely elucidated, the N-terminal region of TRAK family proteins contain a kinesin-interacting domain (Smith et al., 2006) as well as coiled-coil regions that may facilitate homodimerization (Glater et al., 2006; Stowers et al., 2002). Additional N-terminal regions mediate interactions with dynein complexes (Fenton et al., 2021), and the TRAK proteins also contain a central, MIRO-binding domain region (MBD) initially described as consisting of approximately 220 amino acids (MacAskill et al., 2009). More recently (and after these studies described below were completed), the MIRO1-binding domain of TRAK1 was further characterized to ∼40 amino acids sufficient for binding to N-terminal fragments of MIRO1 *in vitro,* and necessary for MIRO1-mediated localization of TRAK1 to mitochondria in cells (Baltrusaitis et al., 2023). TRAK1 has also been shown to activate both kinesin and dynein motility in *in vitro* reconstitution assays (Canty et al., 2023). These findings have shaped the perception of MIRO as a required component of mitochondrial localization (Guo et al., 2005). However, there is evidence that TRAK may interact with mitochondria independently of its association with MIRO. Recent studies have shown that when TRAK is expressed in cells lacking endogenous MIRO, TRAK proteins still localize to mitochondria (Lopez-Domenech et al., 2018), suggesting alternative mechanisms for TRAK/MOM interaction and potentially a distinct role for MIRO in MOM/motor interactions beyond serving as an anchor.

In considering the mechanism of interaction between mitochondria and MYO19, Bocanegra and colleagues proposed a model where MIRO proteins serve recruitment MYO19 through fast-exchanging, weaker-binding interactions, and that MIRO-independent binding facilitates longer-term, stronger-binding docking between the MOM and MYO19 (Bocanegra et al., 2019). The discovery of MIRO-independent TRAK binding to the MOM by López-Doménech and colleagues led us to the hypothesis that TRAK/MIRO interactions may also serve additional roles beyond association, and that a MIRO-independent MOM binding region could be identified in the TRAK proteins. If the mechanism of TRAK/MOM interaction was similar to the mechanisms observed for MYO19/MOM interactions, then MIRO-independent mechanisms of TRAK binding to the MOM may have different exchange kinetics than MIRO-dependent mechanisms. We combined end-point quantitative microscopy approaches paired with fluorescence recovery after photobleaching (FRAP) analyses of dynamic behaviors to investigate the properties of TRAK/MOM/MIRO interactions. Those observations led us to a model for MIRO function where faster-exchanging TRAK/MIRO interactions facilitate the establishment of MIRO-independent TRAK/MOM binding. We were able to identify the MIRO-independent MOM binding domain to the C-terminus of both TRAK1 and TRAK2, and steady-state kinetic data lends further support to a model where MIRO proteins serve in the faster-dissociating recruitment of proteins to the MOM, while MIRO-independent binding mediates slower-dissociating interactions with the mitochondria.

## Results and Discussion

### Ectopic MIRO2 expression enhances TRAK binding to mitochondria

HeLa cells were used to quantify the impact of increased MIRO expression on the association of GFP-TRAK constructs with mitochondria via mito-cyto ratio analysis. Transfection conditions were chosen so that levels of ectopic expression were not saturating so as not to mask mitochondrial localization. GFP-tagged versions of either TRAK showed enriched mitochondrial localization relative to cytosolic localization (MCR>1), and ectopic mchr-MIRO expression enhanced the localization of GFP-TRAK proteins to mitochondria (Figure 1B and 1D). Ectopic expression of mchr-MIRO1 also enhanced GFP-TRAK^MBD^ localization to mitochondria for both TRAK proteins, while expression of mchr-MIRO2 increased the MCR for cells expressing GFP-TRAK1^MBD^. There was little evidence of mitochondrial localization of the TRAK^MBD^ constructs in the absence of ectopic mchr-MIRO (Figure 1C and 1E). HeLa cells endogenously express both MIRO1 and MIRO2 (Oeding et al., 2018), so it may be initially surprising that TRAK^MBD^ constructs did not localize strongly to mitochondria when expressed in the absence of exogenous mchr-MIRO. As the interaction between TRAK^MBD^ and MIRO may be a potentially low-affinity interaction, these results suggest that the endogenous levels of MIRO expression in HeLa cells are such that GFP-TRAK^MBD^ binding is not easily detectable by our method.

**Figure 1:**
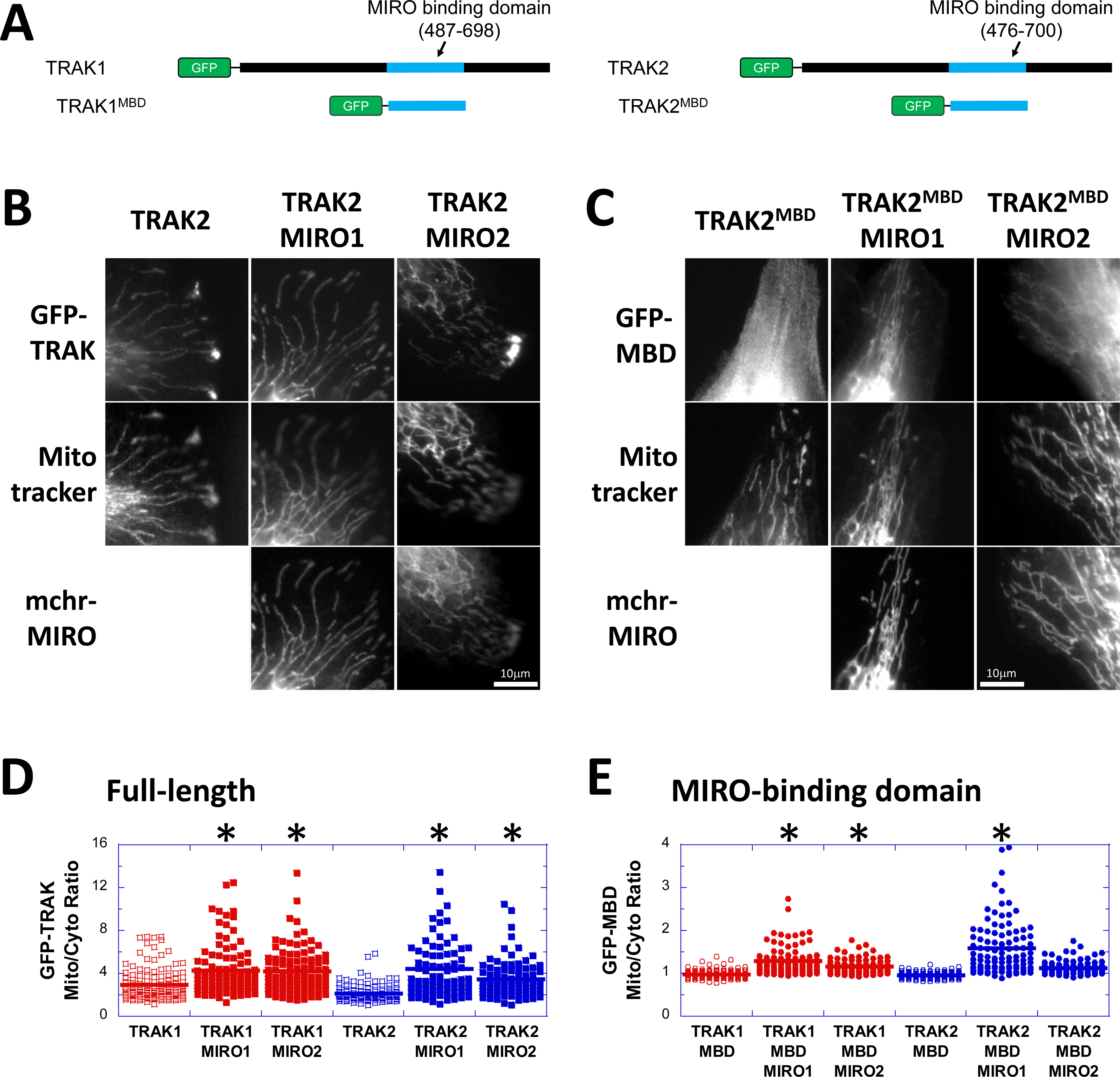
Quantitative image analysis of TRAK1 & TRAK2 localization shows that ectopic expression of mchr-MIRO2 enhances TRAK-binding to mitochondria in HeLa cells. (A) GFP domains were attached to full length TRAK and the TRAK^MBD^, for both TRAK1 and TRAK2. (B) Strong mitochondrial localization was the most common pattern observed in cells expressing full length GFP-TRAK, even in the absence of mchr-MIRO coexpression. (C) GFP-TRAK2^MBD^ remained primarily cytosolic unless coexpressed with an mchr-MIRO. (D) Full length GFP-TRAK1 or GFP-TRAK2 localized more strongly to mitochondria than to the cytoplasm as indicated by a MCR>1, and enrichment was enhanced by ectopic expression mchr-MIRO1 or mchr-MIRO2 (*p<0.0001, Dunnett’s test compared to the corresponding GFP-TRAK construct expressed alone, n ≥ 83 cells per condition). (E) GFP-TRAK^MBD^ constructs only showed mitochondrial enrichment when expressed in combination with mchr-MIRO (*p<0.0001, Dunnett’s test compared to the corresponding GFP-TRAK^MBD^ construct expressed alone, n ≥ 91 cells per condition). The patterns observed for GFP-TRAK2-expressing cells in panels (B) and (C) are also representative of the patterns observed for GFP-TRAK1-expressing cells under the same conditions. Scale bar, 10 μm.

As the full-length GFP-TRAK proteins displayed easily observable mitochondrial localization in the absence of ectopic MIRO expression (MCR ∼2, Figure 1D), and the MIRO-enhancement of TRAK^MBD^ localization was not as pronounced (MCR ∼1.3, Figure 1E), we surmised that the mitochondrial localization displayed by the full-length GFP-TRAK constructs was not solely due to TRAK-MIRO interactions. López-Doménech and colleagues recently generated mouse embryonic fibroblasts lacking MIRO1 or MIRO2 expression--MEF^dko^ cells. MIRO proteins are often described as the linkage between the MOM and TRAK proteins (and often illustrated this way, Fenton et al., 2020; Kruppa and Buss, 2021; Panchal and Tiwari, 2021). However, when MEF^dko^ cells were characterized, TRAK proteins could be detected in mitochondria-enriched cellular fractions and GFP-TRAK proteins localized with mitochondrial markers, lending further support to the presence of a MIRO-independent MOM-binding domain in TRAKs (Lopez-Domenech et al., 2018).

To further investigate how the presence or absence of MIRO proteins influenced TRAK localization to the MOM, we repeated MCR analysis for GFP-TRAK constructs in MEF^dko^ cells. In this cellular context, full-length GFP-TRAK constructs still display an MCR>1, and expression of mchr-MIRO1 increased the MCR for GFP-TRAK1 and TRAK2. mchr-MIRO2 expression also increased the MCR for GFP-TRAK2 (Figure 2A). Expression of mchr-MIRO was required for mitochondrial localization of GFP-TRAK^MBD^ constructs (Figure 2B), with mchr-MIRO2 expression enhancing mitochondrial localization for the MBD domains of both TRAKs while mchr-MIRO1 expression enhanced GFP-TRAK2^MBD^ localization.

**Figure 2:**
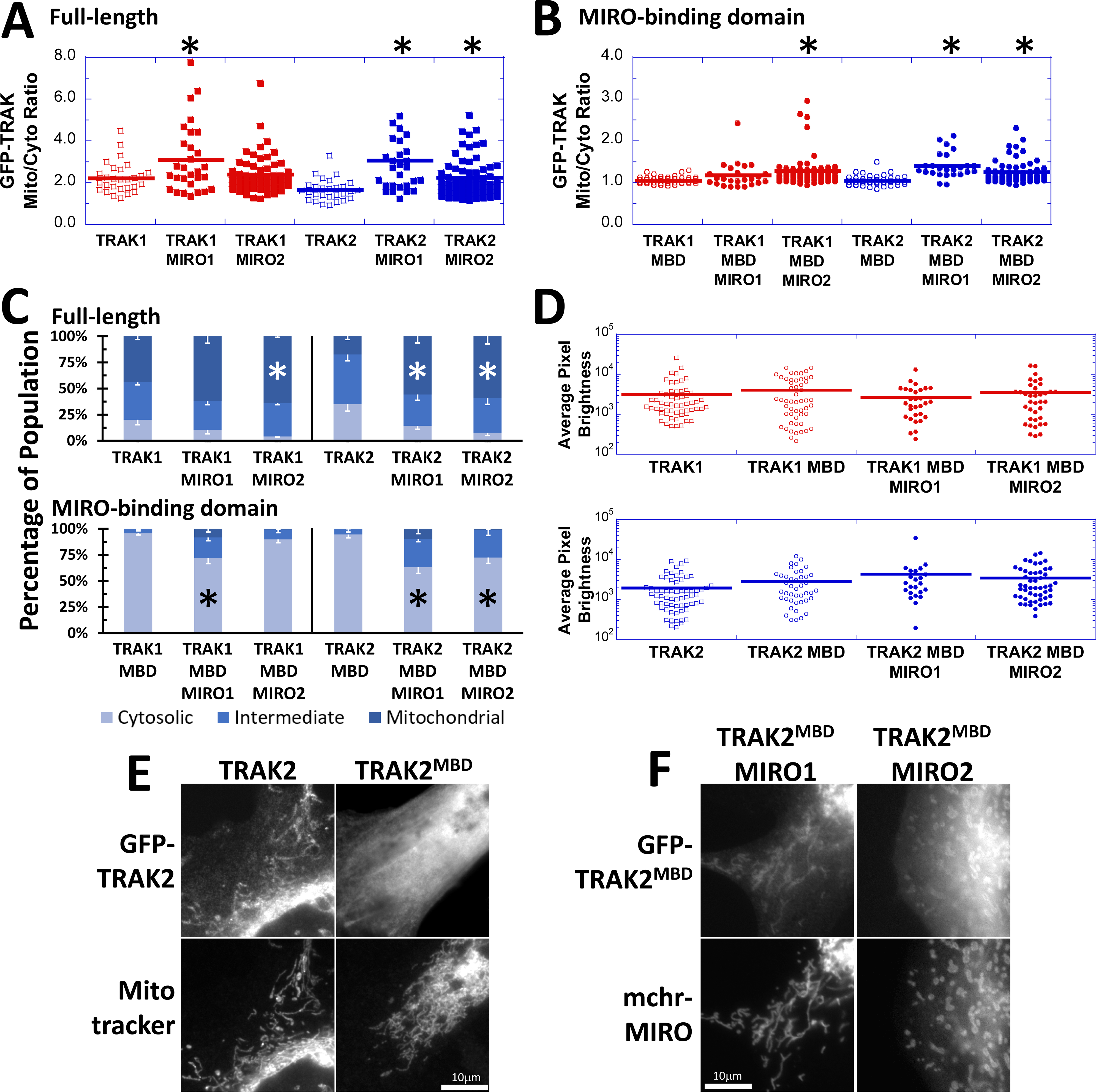
Ectopic mchr-MIRO2 expression enhanced GFP-TRAK localization to mitochondria in MEF^dko^ cells. (A) Full length GFP-TRAK1 or GFP-TRAK2 constructs displayed enhanced fluorescence on mitochondria relative to cytoplasm, even in the absence of MIRO expression (MCR > 1). In some instances, coexpression of an mchr-MIRO increased the relative brightness of the GFP-TRAK fluorescence on mitochondria as compared to the cytoplasm (*p<0.04, Dunnett’s test compared to the corresponding GFP-TRAK construct expressed alone, n >= 28 cells per condition). (B) GFP-TRAK^MBD^ constructs only displayed enhanced mitochondria fluorescence when coexpressed with an mchr-MIRO (*p<0.0001, Dunnett’s test compared to the corresponding GFP-TRAK^MBD^ construct expressed alone, n >= 25 cells per condition). (C) When assessed for the fraction of cells displaying mitochondrial localization of a GFP-TRAK construct, coexpression of mchr-MIRO decreased the percentage of cells showing primarily cytosolic localization of GFP-TRAK1 and GFP-TRAK2 in some instances (*p<0.02, Dunnett’s test compared to the corresponding GFP-TRAK construct expressed alone, n = 4 or 5 independent experiments per condition, error bars represent SEM). GFP-TRAK^MBD^ constructs displayed a larger increase in the fraction of cells with an intermediate or mitochondrial phenotype when coexpressed with an mchr-MIRO (*p<0.02, Dunnett’s test compared to the corresponding GFP-TRAK^MBD^ construct expressed alone, n = 5 independent experiments per condition, error bars represent SEM). (D) It is unlikely that phenotypic differences observed with respect to MCR for individual cells and the fraction of the population displaying cytosolic localization for GFP-TRAK constructs are due to differences in protein expression such that mitochondria-localized GFP-is masked by high cytoplasmic signal. All constructs displayed similar distributions of protein expression and average pixel brightness was not different for either TRAK1 or TRAK2 (Dunnett’s test compared to the corresponding GFP-TRAK^MBD^ condition, n >= 22 individual cells per condition). Panels (E) and (F) show representative images of the common patterns observed for mitochondrial localization in cells expressing GFP-TRAK2 constructs. These images are also representative of the patterns observed for GFP-TRAK1-expressing cells under the same conditions. Scale bar, 10 μm.

To further characterize the prevalence of the observed phenotypes, population phenotype analysis was performed in MEF^dko^ cells expressing different combinations of GFP-TRAK and mchr-MIRO constructs. Ectopic expression of mchr-MIRO2 reduced the fraction of cells displaying a cytosolic GFP distribution in cells coexpressing GFP-TRAK1, GFP-TRAK2, or GFP-TRAK2^MBD^, but not for GFP-TRAK1^MBD^. Expression of mchr-MIRO1 increased the fraction of cells with intermediate or mitochondrial localization when coexpressed with full-length GFP-TRAK2 or with either GFP-TRAK^MBD^ construct (Figure 2C). Even though care was taken in limiting ectopic expression of the GFP-tagged constructs, one possible explanation for the differences in MCR and the fraction of cells with strong mitochondrial localization would be that overexpression levels were so great that any mitochondrial localization was masked by high levels of cytosolic fluor due to saturation of the available binding sites. We calculated the average GFP-pixel brightness for cells expressing the TRAK constructs alone, or in combination with mchr-MIRO. For all cell types the pixel brightness ranged similarly, and the average brightness was not significantly different between GFP-TRAK and GFP-TRAK^MBD^ whether the MBD construct was expressed alone, or in combination with mchr-MIRO (Figure 2D). These data support our interpretation that the conditions where MCR was enhanced and where the population of cells displayed a higher fraction of cells with a mitochondria-localization phenotype is due to a redistribution of GFP-TRAK signal to the mitochondria as a result of mitochondrial binding (Figure 2E and 2F).

The differential impact of MIRO expression on GFP-TRAK1 and GFP-TRAK2 constructs in MCR analysis and in population analysis could indicate differences in affinity between the two MIRO and the two TRAK isoforms. These differences could be indicative of differences in cellular roles between TRAK1 and TRAK2 as well as between MIRO1 and MIRO2 (Lopez-Domenech et al., 2018; van Spronsen et al., 2013). Taken together, these findings confirm a MIRO-independent mechanism of TRAK localization to the MOM.

### MIRO-independent mitochondrial localization is encoded by the C-terminus of TRAK proteins

To investigate which region of TRAK facilitates MIRO-independent binding, we generated GFP-TRAK truncations encoding for the N-terminal region before the MBD and the C-terminal region after the MBD for both TRAK1 and TRAK2 (Figure 3A and S1). Previous studies had identified a role for the C-terminus of Milton, the *Drosophila* ortholog of TRAK proteins, in mitochondrial localization. However, since those constructs were expressed in cells with endogenous MIRO expression (Glater et al., 2006), even though these constructs did not contain the putative MIRO-binding domain, a second mechanism of MIRO-dependent binding could not be ruled out. When expressed in MEF^dko^ cells, neither of the TRAK^N-term^ constructs showed enhanced mitochondrial localization, while both TRAK^C-term^ constructs did display an MCR >1 (Fig. 3B and 3C). Population phenotype analyses also revealed similar behavior--cells expressing the GFP-TRAK^N-term^ displayed primarily cytosolic GFP fluorescence while cells expressing the TRAK^C-term^ had a greater fraction of intermediate and mitochondrial GFP localization, further supporting the presence of a MIRO-independent MOM association domain encoded by the C-terminus of TRAK (Figure 3D and 3E).

**Figure 3:**
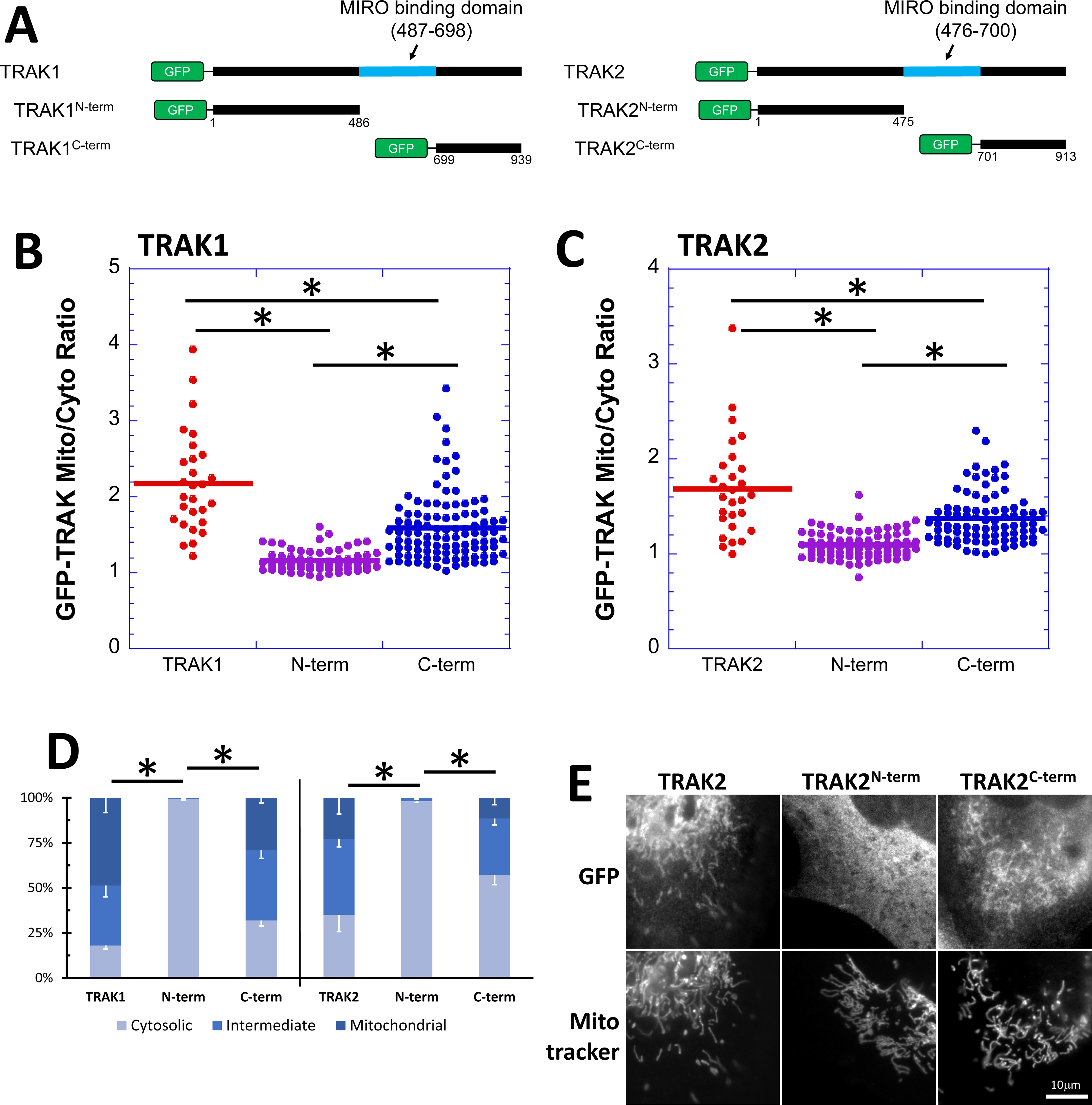
The MIRO-independent mitochondrial localization domain of TRAK is encoded by the amino acids c-terminal of the MIRO-binding domain. (A) GFP domains were attached to full length TRAK as well as N-terminal and C-terminal fragments for both TRAK1 and TRAK2. (B) GFP-TRAK1^C-term^ expressed in MEF^dko^ cells displays brighter fluorescence in mitochondrial regions as compared to the cytosol regions, but to a lower degree than GFP-TRAK1. GFP-TRAK1^N-term^ has an MCR of ∼1 and does not show mitochondrial localization (*p<0.0001, Dunnett’s test compared to full-length, n ≥ 28 cells). (C) Similar localization patterns were observed for GFP-TRAK2 (*p<0.0001, Dunnett’s test compared to full-length, n ≥ 28 cells). (D) The fraction of cells displaying primarily cytoplasmic fluorescence was smaller for full-length GFP-TRAK or the GFP-TRAK^C-term^ constructs as compared to the GFP-TRAK^N-term^ constructs (*p>0.002, Dunnett’s test compared to the corresponding GFP-TRAK^N-term^ construct, n ≥ 3 independent experiments per condition, error bars represent SEM). Panel (E) shows representative patterns observed in cells expressing GFP-TRAK2 constructs, and are also representative of the patterns observed in cells expressing GFP-TRAK1 constructs. Scale bar, 10 μm.

We attempted to identify the minimal MOM association domain (TRAK^min-C^) by using MemBrain3.1 (Feng et al., 2022), which was employed by Shneyer and colleagues in identifying the MIRO-independent MOM interacting domain found in MYO19 (Shneyer et al., 2016). The TRAK^min-C^ sequences predicted by MemBrain were also predicted by AlphaFold (Jumper et al., 2021; Varadi et al., 2022) to contain alpha helical regions (Figure S2). However, neither GFP-TRAK1^min-C^ nor GFP-TRAK2^min-C^ constructs localized to mitochondria in MEF^dko^ cells, whether the GFP-tag was at the N-terminus of the truncation or the C-terminus of the truncation (data not shown). The predictive power of the informatic approaches were insufficient in this instance. The potential TRAK binding partners mediating this interaction includes the mitofusins (Lee et al., 2018; Misko et al., 2010), while other potential binding partners have been identified via proteomic analyses (Oughtred et al., 2021).

### The differences in the FRAP kinetics of the TRAK domains responsible for mitochondrial binding indicate differences in association with the MOM

Fluorescence recovery after photobleaching (FRAP) was performed to determine the kinetic properties of TRAK/MOM interactions mediated through MIRO-dependent or MIRO-independent mechanisms. While GFP-TRAK transfected cells had a range of cytosolic signal, all cells chosen for FRAP analysis had an MCR>1 and strong mitochondrial localization that was readily apparent by eye. Fluorescence recovery could be a result of equilibrium exchange of photobleached material on mitochondrial structures with unbleached cytoplasmic pools. If a subregion of reticular mitochondrial network was bleached, recovery could potentially occur through exchange of material from within the mitochondrial network. Additionally, unbleached mitochondrial particles could translocate from other regions of the cell and fuse with bleached mitochondrial networks. Isolated mitochondria without obvious connectivity to the rest of the mitochondrial network displayed fluorescence recovery (Figure S3A), indicating exchange with cytoplasmic pools. Depending on the GFP construct, photobleached regions within more extensive mitochondrial networks could be seen to recover from the margins of the bleached region towards the center (Figure S3B). Rarely we would observe movement and fusion of unbleached mitochondria with bleached mitochondrial networks, resulting in a rapid decrease in fluorescence of the “donor” organelle (Figure S3C). Cytoplasmic recovery and network recovery were observed most often, and kinetic data were fit to a double-exponential rise to account for FRAP recovery resulting from the cooccurrence of these two distinct mechanisms.

Full-length GFP-TRAK1 and GFP-TRAK1^C-term^ expressed in MEF^dko^ cells had 6-to 14-fold slower rates of exchange compared to that of GFP-TRAK1^MBD^ expressed in combination with either mchr-MIRO, as measured by t_1/2_ (Fig. 4A). Similar results were observed with GFP-TRAK2 constructs (Fig. 4B), where GFP-TRAK2^MBD^ constructs expressed in combination with mchr-MIRO displayed a 7-fold faster t_1/2_ compared to GFP-TRAK2 (Figure 4 and S4B). GFP-TRAK^C-term^ constructs displayed FRAP kinetics at an intermediate level between the TRAK^MBD^ and the full-length construct.

**Figure 4:**
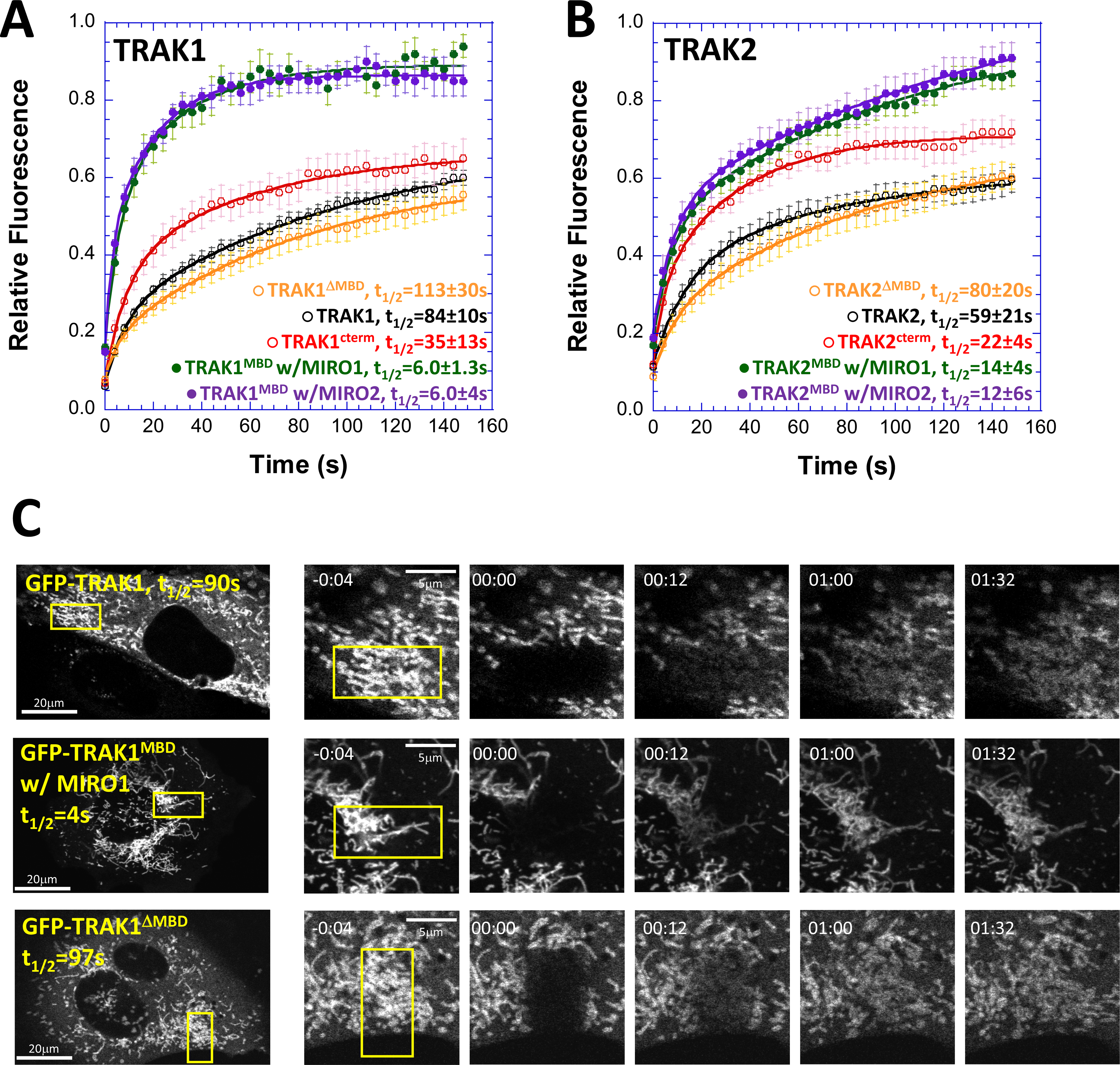
FRAP analysis reveals that GFP-TRAK^MBD^/mchr-MIRO exchange kinetics are faster than MIRO-independent exchange kinetics for other GFP-TRAK constructs. GFP-tagged TRAK 1 (A) and TRAK2 (B) constructs were expressed in MEF^dko^ cells, were then analyzed for fluorescence recovery after photobleaching in cells showing strong mitochondrial localization. Constructs containing the C-terminal domain were capable of mitochondrial localization independent of MIRO expression and had significantly larger t_1/2_, compared to mitochondria-localized GFP-TRAK^MBD^ in the presence of mchr-MIRO1 or mchr-MIRO2. Markers represent the mean ± SEM at each time point, and the curves are a double-exponential rise calculated from the averaged data, n>= 6. See Figure S3 for a summary of the analyses for all conditions. (C) Representative imaging series for FRAP samples. Fluorescence returned more rapidly to mitochondria for GFP-TRAK1^MBD^ in the presence of mchr-MIRO1 than it did to GFP-TRAK1 or GFP-TRAK1^ΔMBD^. The patterns observed in these images are representative of the patterns observed in cells expressing different combinations of constructs but with similar FRAP kinetics. The photobleached region is indicated by the yellow box. Time is indicated in minutes:seconds.

The differences in FRAP kinetics between the full-length and C-terminal constructs parallel what was observed with end-point measures (MCR and population analysis). The enhancements in mitochondrial localization (Figure 3) and in FRAP t_1/2_ observed with the C-terminal constructs was less robust than what was seen with the full-length constructs. It is possible that regions in the N-terminus insufficient for mitochondrial localization also serve to enhance the properties encoded by the C-terminus. Multimerization through coiled-coil domains found in the N-terminus (Glater et al., 2006; Stowers et al., 2002) would increase the valence of MOM association domains in any one protein assembly, influencing its MOM interaction kinetics. To investigate this question, we conducted FRAP analysis on GFP-TRAK constructs lacking the MIRO-binding domain (GFP-TRAK^ΔMBD^, see Figure S1 for a schematic). The observed t_1/2_ for these constructs were slower than the t_1/2_ for the corresponding GFP-TRAK^C-term^ construct and similar in order of magnitude to the t_1/2_ observed with the full-length constructs, supporting the hypothesis that N-terminal sequences are necessary for exchange kinetics similar to that of the full-length protein, even though the N-terminal sequence is insufficient to encode for MOM localization (Figure 4 and S4). Just as multimerization of motor proteins influences the time-fraction that the proteins are associated with their cytoskeletal filament (Howard, 2001), dimerization of TRAK through the N-terminal coiled-coil domains would enhance the likelihood of a protein assembly binding to the MOM as the multimer contains more potential binding sites than the monomer.

To establish frames of reference for other putative components of motor complexes on the MOM in a similar cellular context, we completed FRAP analyses for GFP-MIRO2, and for the regions of MYO19 capable of MIRO-dependent (GFP-MYO19^MBD^ expressed with mchr-MIRO2) and MIRO-independent (GFP-MYO19^mem^) mitochondrial association in MEF^dko^ cells (Figure S4). As we had previously reported in HeLa cells (Bocanegra et al., 2019), the t_1/2_ for GFP-MYO19^mem^ was larger than that of GFP-MYO19^MBD^/mchr-MIRO2, indicating faster exchange for MIRO-mediated association. It is worth noting that the t_1/2_ of the GFP-MYO19^MBD^/mchr-MIRO2 interaction was slower in MEF^dko^ cells than what was previously reported in HeLa cells. One possible explanation for this difference is the cellular context. HeLa cells express MIRO1 and MIRO2, and the sum of endogenous and exogenous binding sites for GFP-MYO19^MBD^ could result in increased opportunities for exchange and an apparent faster recovery of fluorescence. Additionally, the t_1/2_ for GFP-MIRO2 was of similar magnitude to that of GFP-MYO19^mem^ and slower than any of the GFP-TRAK constructs containing MIRO-independent MOM binding ability (Figure S4), suggesting that the MIRO-independent mechanisms binding to the MOM may differ mechanistically between MYO19 and TRAK. To get a sense of the dissociation behavior of the MIRO-dependent and MIRO-independent mechanisms that TRAK1 employs to bind to mitochondria, we examined the nonequilibrium dissociation of GFP constructs via digitonin-induced permeabilization activated reduction in fluorescence (Bocanegra et al., 2019; Hawthorne et al., 2016; Singh et al., 2016), and PARF analysis indicated that the off-rate kinetics for GFP-TRAK1 and GFP-TRAK1^C-term^ were approximately 10-fold slower than the t_1/2_ observed for GFP-TRAK1^MBD^ expressed in combination with either mhr-MIRO1 or mchr-MIRO2 (Figure S5).

Taken together, these analyses suggest that GFP-TRAK^C-term^ interactions with the MOM have a stronger association than the GFP-TRAK^MBD^ interactions with mchr-MIRO. As the kinetics of full-length GFP-TRAK are also significantly slower than those between GFP-TRAK^MBD^ and mchr-MIRO, these data may indicate that the TRAK^MBD^ interaction with MIRO facilitates a weaker, faster-exchanging recruitment of TRAK to the MOM while TRAK^C-term^ is partially responsible for slower turnover and stronger interactions between TRAK and the MOM.

### Kinetic considerations influence our understanding of MIRO function in the recruitment of cytoskeletal motors to mitochondria

Current models suggest that MIRO “anchors” TRAK proteins to the mitochondrial membrane (Kruppa and Buss, 2021). This interpretation is likely a consequence of experimental design--many of the previous experimental examinations of MIRO/TRAK interactions were carried out as end-point assays. Cells were made to express (or not express) a set of proteins under particular conditions, and then “fixed in time” for analysis. These analyses included assaying for binding via immunoprecipitation, or for co-occurrence via qualitative fluorescence microscopy. Previous experimental approaches do not address time as key component required for full understanding of biologically relevant interactions. Additionally, solely qualitative imaging approaches make it more difficult to validate and compare results between experimental conditions (Wait et al., 2020).

Kinetic descriptors are crucial to informing a more complete biological understanding (Pollard and De La Cruz, 2013), and we believe this to be the first reported kinetic analysis of TRAK/MIRO interactions. In taking a quantitative and kinetic approach to our experimental design, we were able to identify relevant differences in the biochemical behavior of specific segments of the TRAK protein sequence with respect to MOM association. For any of these observations to be valid, we had to verify that observed phenotypic differences were unlikely to be due to differences in levels of overexpression (Figure 2D). Population and MCR analyses in MEF^dko^ cells lacking MIRO expression enabled us to further characterize the ability of TRAK proteins to localize to mitochondria independent of MIRO (Figure 2) and that the C-terminus of TRAK encoded for this property (Figure 3). Quantitative fixed-cell endpoint assays supported the idea that the TRAK^C-term^/MOM interaction likely has different properties than that of the TRAK^MBD^/MIRO interaction, and it is worth noting that the TRAK^C-term^ does not overlap with the sequences involved in dynein or kinesin interactions (Fenton et al., 2021). We were able to demonstrate differences in binding kinetics between MIRO-dependent and MIRO-independent interactions with the MOM only by employing experimental approaches that allowed quantification of dynamic molecular processes (Figure 4 and Figure S2). Although absolute concentrations of relevant components cannot be known in these assays, the similarities in relative levels of expression (Figure 2D) allow us to make comparisons between our experimental conditions.

### A model for the role of MIRO in motor protein/mitochondria interactions

The application of kinetic approaches led us to a model of MIRO function where the role of MIRO could be as a low-affinity acceptor of motor proteins and their adaptors, while other mechanisms are responsible for higher-affinity, longer-lived attachments (Figure 5). Our observations with TRAK proteins parallel what we had previously reported for MYO19, where kinetic analysis of MYO19/MIRO interactions first revealed that MIRO may provide fast-exchanging binding sites on the MOM for proteins that also contained a second, slower exchanging MOM association domain (Bocanegra et al., 2019). In this way, active MIRO could enhance recruitment of cytoskeletal motors and their adaptors to the MOM, potentially increasing their local concentration. Higher local concentrations of MIRO binding partners could support formation of MIRO-independent interactions with the MOM which might be tighter-binding, but might also be lower probability events. This initial recruitment activity could potentially be regulated by nucleotide state-dependent interactions between the N-terminal GTPase domain of MIRO and its binding partners (Bocanegra et al., 2019; Fransson et al., 2006). If MIRO were in an inactive state, recruitment would be halted, but it would not lead to dissociation of motor complexes attached to the MOM through MIRO-independent mechanisms.

**Figure 5:**
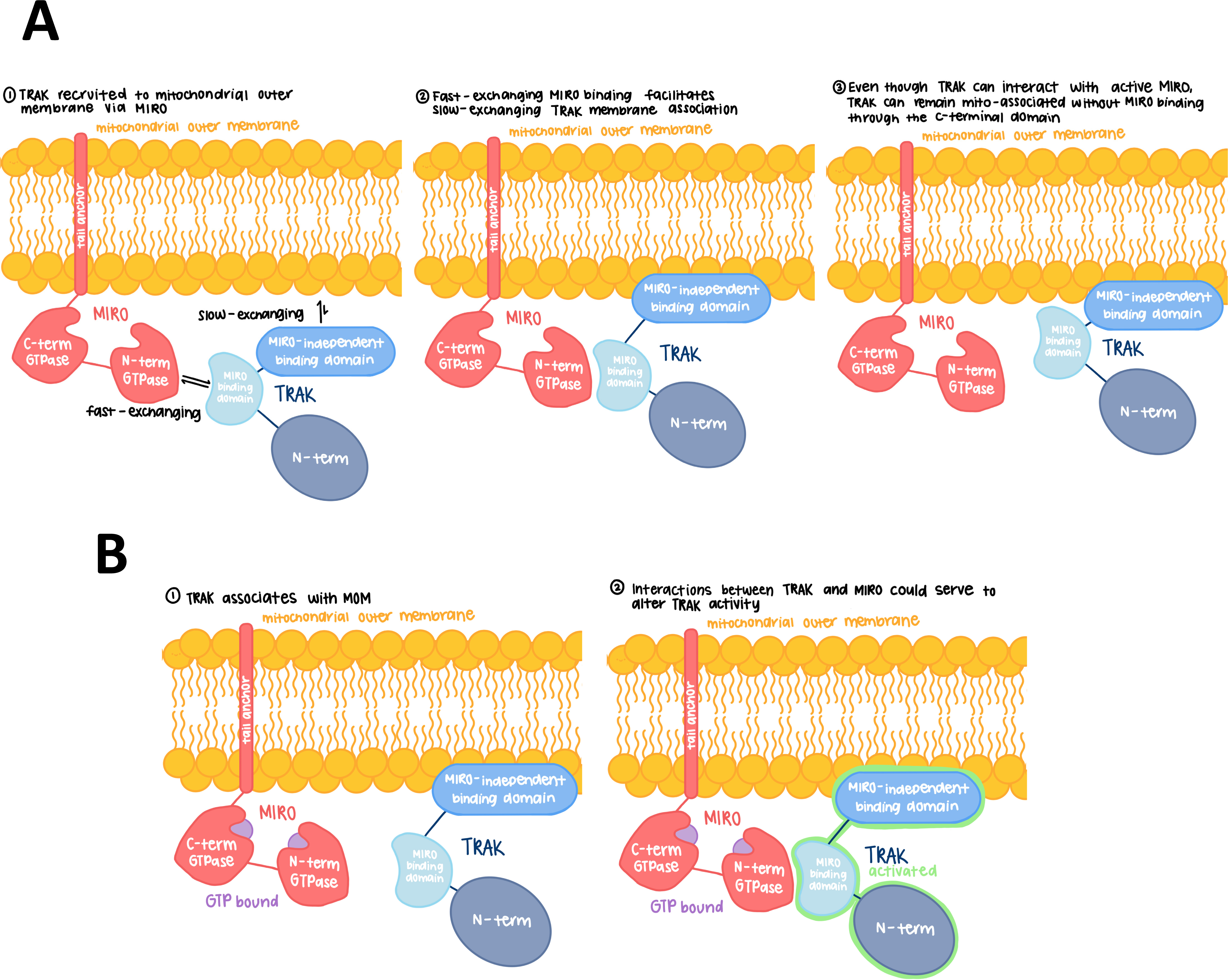
A kinetic model for MIRO interactions with molecular motor assemblies on the mitochondrial outer membrane. Our data indicate that TRAK is capable of MIRO-independent mitochondrial binding, and that MIRO-independent binding has slower turnover kinetics, indicating a tighter interaction than MIRO-dependent binding. (A) Therefore, MIRO may be recruiting TRAK and their associated MT motors to mitochondria via lower affinity interactions, while tighter binding is mediated by the C-terminal domain. As a similar mechanism exists for MYO19, MIRO’s function may be as an initial recruiter of a number of different proteins to the mitochondrial outer membrane, even if their mechanisms of MOM binding are different. (B) Interactions between active MIRO and its MOM-associated binding partners could regulate the activity of those partners.

This idea is supported by the observations by Oeding and colleagues as well as López-Doménech and colleagues (Lopez-Domenech et al., 2018; Oeding et al., 2018). Both groups observed that in cells lacking MIRO proteins, MYO19 levels decreased significantly. Their data also supported the interpretation that without MIRO proteins to assist in the insertion of MYO19 with the MOM, MYO19 remains cytosolic: accessible to ubiquitination and proteasome-mediated degradation.

It is curious that mitochondria-associated TRAK protein persist in the absence of MIRO expression even though a similar pattern for the kinetics of MIRO-dependent (faster) versus MIRO-independent (slower) MOM association was observed. This could be explained by the difference in the t_1/2_ order of magnitude between the MIRO-independent interactions for TRAK and MYO19--the MIRO-independent TRAK/MOM interaction has a ∼5-fold faster t_1/2_ than the MYO19/MOM interaction (Figure S4). MIRO-independent TRAK binding to the MOM may still happen at rates sufficiently quick to generate a population of mitochondria-bound TRAK protected from proteolysis, while the MIRO-independent MOM binding for MYO19 might be so infrequent that a larger fraction of the cellular MYO19 pool is susceptible to proteasome activity in the absence of MIRO. Such proteolytic susceptibility results in little to no MYO19 available to associate with the MOM.

If MIRO proteins serve different roles beyond long-term MOM association, then the role of MIRO activation on the availability of the MOM for motor attachment may need to be evaluated more carefully. The current model of MIRO activity is that MIRO may be active and bound to cytoskeletal motors in one nucleotide state, while, favoring the dissociation of motors in a different nucleotide state (Fransson et al., 2003; MacAskill et al., 2009). Our data support an additional model where the nucleotide-state dependence of MIRO is not simply turning the molecular switch from “motors bound” or “motors unbound.” Instead, MIRO nucleotide state is regulating initial MOM binding. As slower-exchanging, MIRO-independent mechanisms also mediate motor/adaptor binding to the MOM, inactivation of MIRO might not result in dissociation of motors/adaptors from the MOM. Alternatively, inactivation of MIRO could inhibit recruitment of additional proteins. In such a model, a mitochondrial network having only GDP-MIRO would not be devoid of motors. Instead, its ability to recruit additional motors would be impaired. Our data support a mechanism where MIRO is analogous to the maître d’ at a Chez Quis. If Abe Frohman, the “Sausage King of Chicago,” and his two guests are not able to be seated (i.e. MIRO in its “off” state), the restaurant patrons who have already been seated would not leave. They would remain at their tables just as TRAK proteins bound to the MOM via the C-terminal domain would remain on mitochondria even with MIRO in its inactive state.

In addition to facilitating recruitment, MIRO may activate binding partners such as the TRAKs. López-Doménech and colleagues showed that the amount of TRAK1 and TRAK2 isolated in mitochondrial fractions was not different between wildtype MEFs and MEF^dko^ cells.

Similar results were observed for microtubule motors as well (Lopez-Domenech et al., 2018). However, MEF^dko^ cells had a number of mitochondria localization and motility defects. The fraction of cell area containing mitochondria was smaller, mitochondrial occupancy of the cell periphery was decreased, the mitochondrial network was less dynamic (as measured by displacement), and fewer individual mitochondria were observed undergoing tubulin-dependent movements in MEF^dko^ cells. One interpretation of these data is that MIRO/TRAK interactions promote TRAK function through activation (Figure 5B), as MIRO-independent localization does not result in the same positioning or dynamics of mitochondria within the cell.

As MIRO interactions with motors are not the exclusive mechanism of mitochondrial interaction, other mechanisms, such as regulated degradation, would contribute to controlling motor protein persistence on the MOM. Future investigations could involve kinetic assays to determine the role of ubiquitin-mediated proteolysis pathways on MOM-associated motor protein levels, and if the lifetime of TRAK adaptors and motor proteins on the MOM is sensitive to particular cellular conditions, such as glucose starvation, inner mitochondrial membrane potential, or cell-cycle state. Our experiments reveal that the homeostatic mechanisms regulating motor protein adaptor attachment to the mitochondria are partially independent of MIRO. Developing a more complete understanding of the processes regulating motor protein adaptor attachment will be as crucial to understanding mitochondrial dynamics as understanding which interactions modulate motor protein (and adaptor) activation once motors are bound to the organelle.

## Acknowledgements

A significant portion of this work was completed during Spring and Summer 2020, at the beginnings of pandemic shutdowns. We would like to thank University of Richmond School of Arts and Science for funding and supporting undergraduate-driven research in the midst of the COVID pandemic. Christie Lacy’s efforts as Director of Imaging and Microscopy, supporting our work while maintain the Imaging Core Facility were essential for the success of this project—we could not make these discoveries without the Core Facility and its Director. We would also like to thank Dr. Andrew Moore (HHMI-Janelia), Dr. Rebecca Adikes (Siena College), and Dr. Teng-Leong Chew (HHMI-Janelia) for giving their time to run out Zoom Image Analysis Bootcamp, introducing “the TRAK team” to the beauty of image quantitation during the summer of 2020. It opened the door for a bunch of naïve undergrads to be scientists even when we could not be in the lab because of the pandemic.

Hypotheses and experiments in this study were conceived by OAQ and LM. Experiments were performed by LM, and OAQ. Data analysis was completed by LM, KR, HF, CV, and OAQ. Manuscript preparation was completed by LM with assistance from KR and OAQ. This work was funded through University of Richmond Arts & Sciences Summer Fellowships to LM, KR, and CV. HF and OAQ were funded by NIH R15 GM119077 to OAQ. A special thank you to Drs. Josef Kittler and Guillermo López-Doménech (University College London) for the MEF^dko^ cells.

---

**Figure S1:**
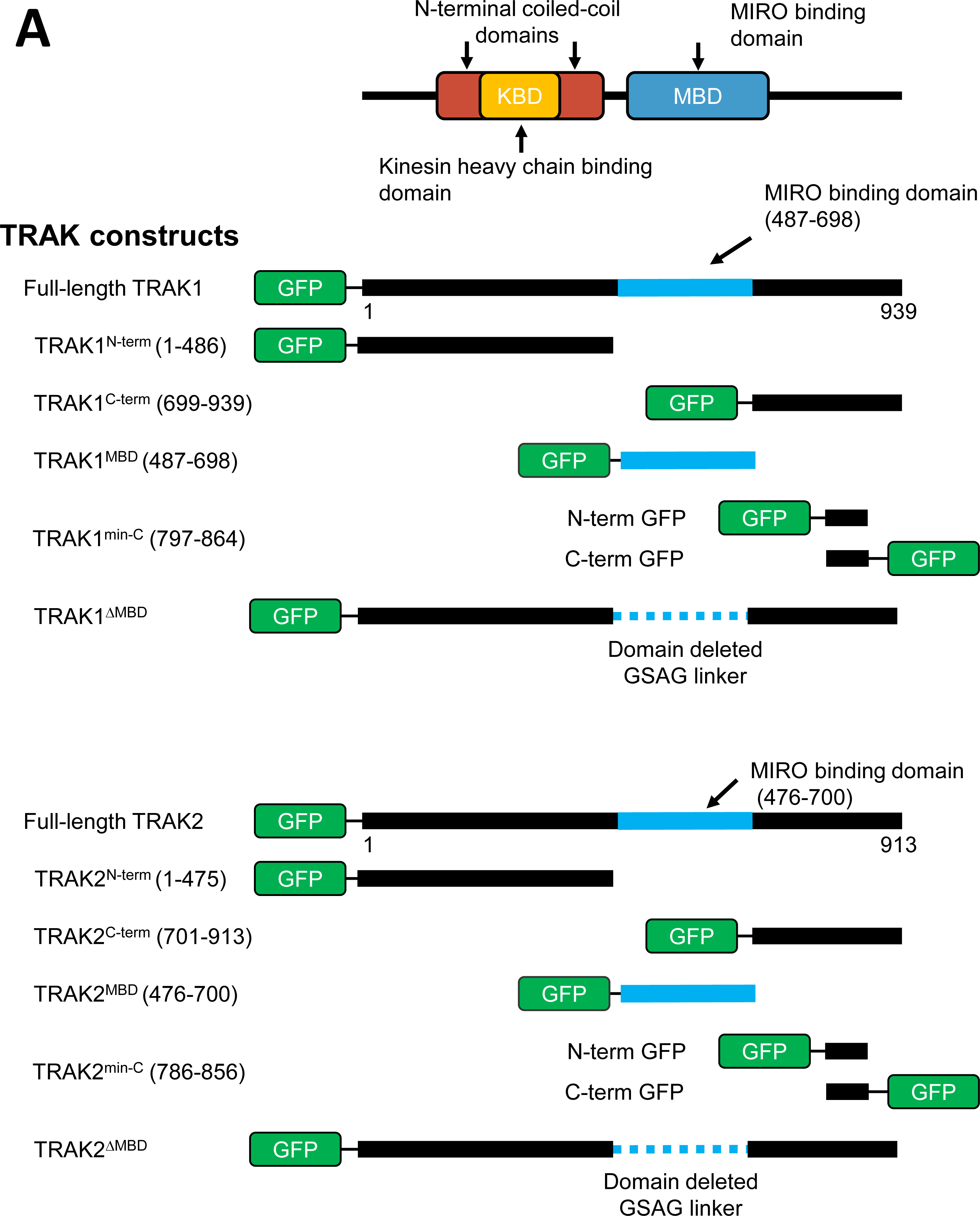
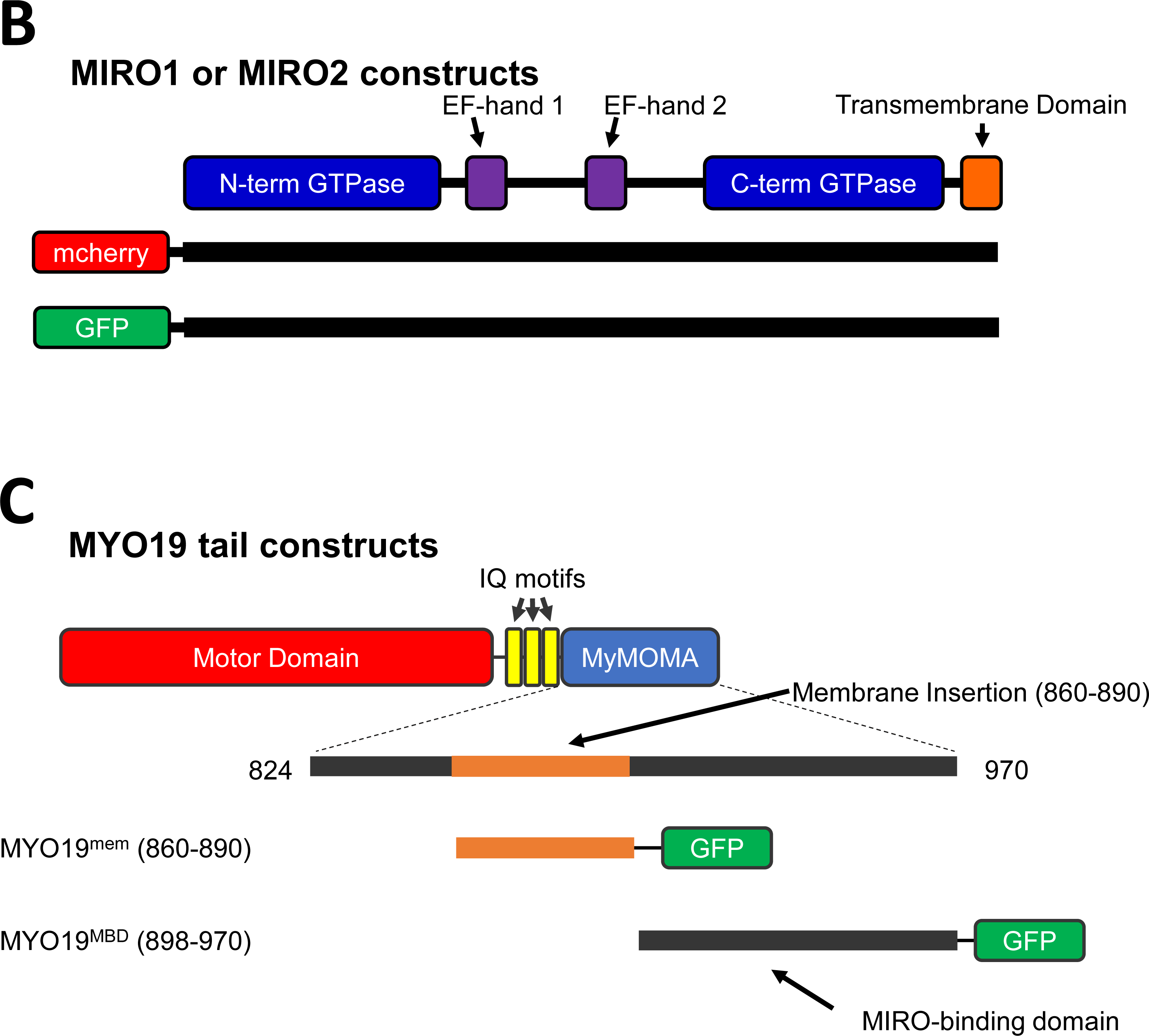
Schematic diagrams of the fluorescent constructs used. GFP-tagged (A) Mouse TRAK1 and TRAK2 constructs were generated from the previously published constructs reported by Birsa et al. (Birsa et al., 2014). In addition to the reported truncations, the amino acid linker between the GFP domain and the TRAK protein was lengthened from SGLRSRAQAS to GEQKLRILQSTSGLRSRAQAS. (B) The mcherry-tagged (Bocanegra et al., 2019) and GFP-tagged (Birsa et al., 2014) human MIRO1 and MIRO2 constructs were previously reported. (C) The human GFP-MYO19^MBD^ (Bocanegra et al., 2019; Oeding et al., 2018) and GFP-MYO19^mem^ (Shneyer et al., 2016) constructs were also previously reported.

**Figure S2:**
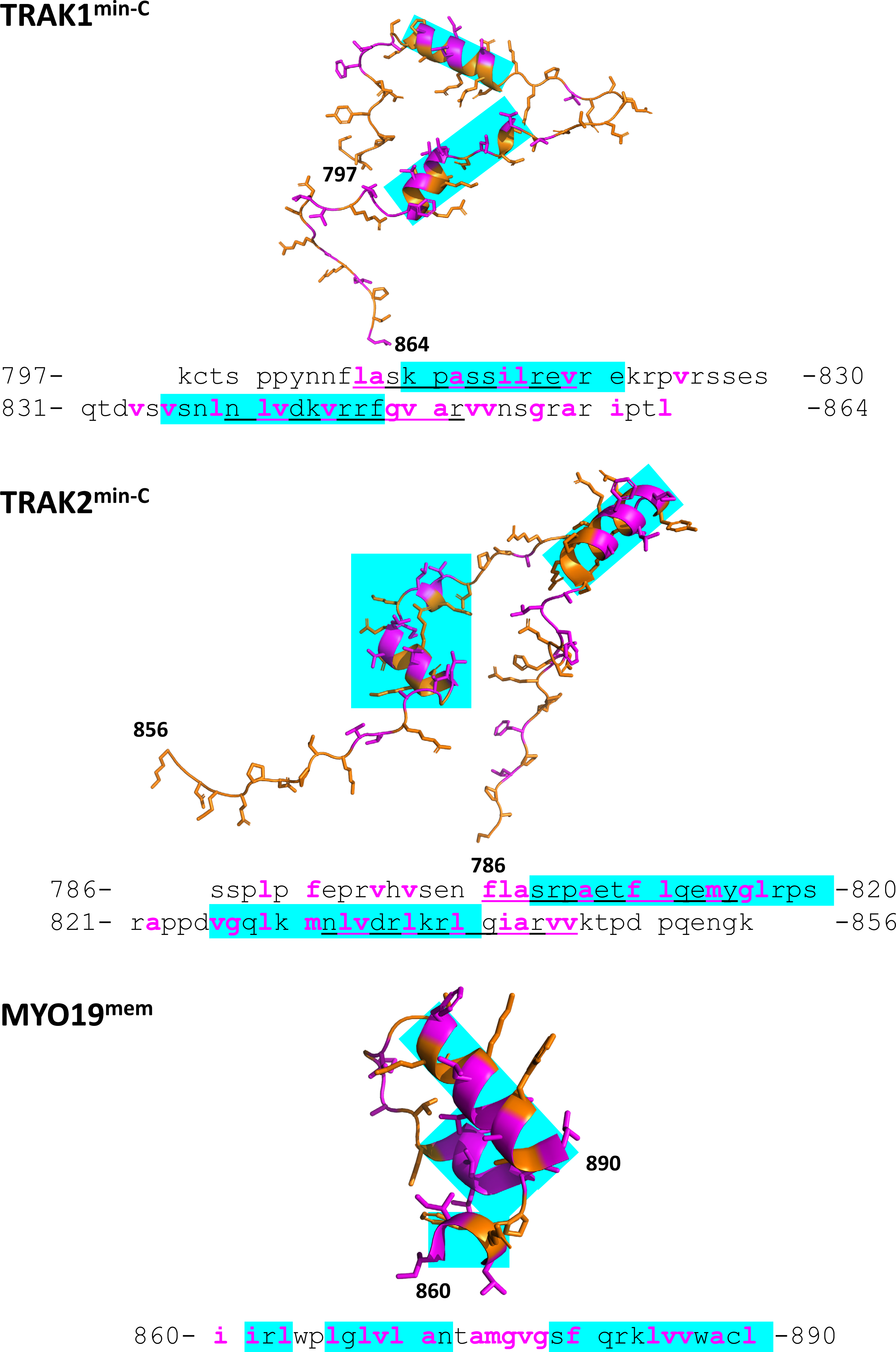
Structural predictions of the minimal domain necessary for MIRO-independent mitochondrial association in the C-terminal regions of TRAK proteins. We used a combination of amphipathic helix prediction via MemBrain 3.1 and structural predictions from AlphaFold to attempt to identify the minimal number of residues encoding for the MIRO-independent mitochondrial association domain of mouse TRAK1 and TRAK2. Magenta letters represent hydrophobic amino acids, cyan highlighting corresponds to AlphaFold helix predictions, and <u>underlines</u> correspond to Membrain 3.1 amphipathic helix predictions (propensity score >0.3). When these truncations were expressed as GFP-tagged proteins in MEF^dko^ cells, neither TRAK1^min-C^ nor TRAK2^min-C^ localized to mitochondria--no matter whether the GFP-tag was N-terminal or C-terminal. For comparison, the MIRO-independent membrane association domain of human MYO19 is also shown, and MYO19^mem^ does localize to mitochondria when expressed as a GFP-construct in MEF^dko^ cells.

**Figure S3:**
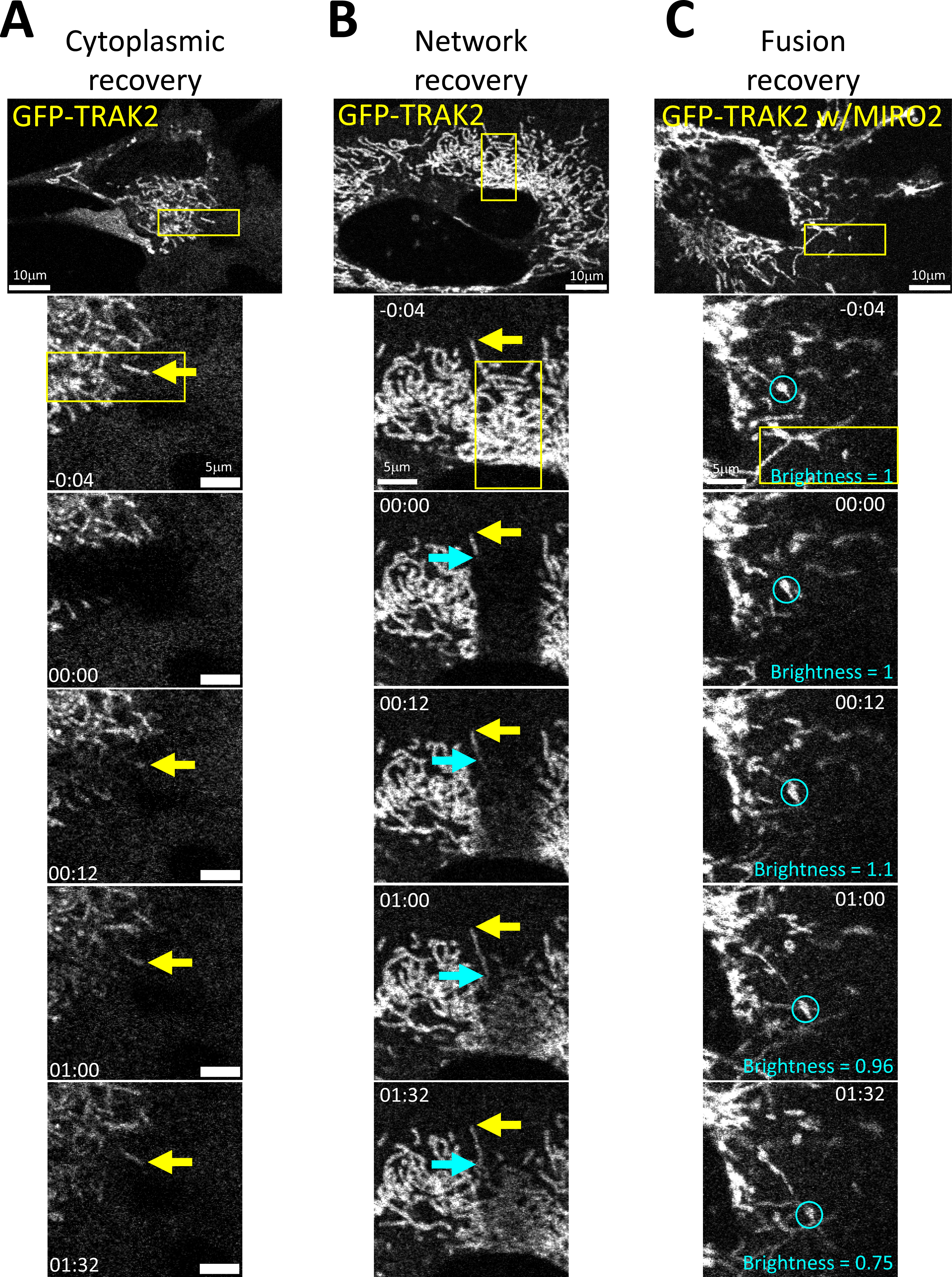
Recovery mechanisms observed in FRAP analysis of GFP-TRAK constructs. Fluorescence recovery after photobleaching results from the exchange of bleached fluors in the region of interest with unbleached fluors from other pools. (A) In isolated mitochondria, we could observe apparent recovery of from cytoplasmic pools (yellow arrow). (B) When a portion of a mitochondrial network was photobleached, we could observe movement of unbleached material into the bleached region (cyan arrow). In this example, the extended mitochondrion recovers fluorescence from the unbleached portion more quickly than the surrounding photobleached regions recover fluorescence. Additionally, the bleached network would often recover from the edges of the bleached region towards the middle. (C) On occasion, we could also observe recovery by movement of unbleached mitochondria into the bleached region (cyan circle). The subsequent fusion led to decreased of mitochondrial fluorescence in the donor organelle with local increased fluorescence in the network to which it had fused. The photobleached region is indicated by the yellow box. Time is indicated in minutes:seconds.

**Figure S4:**
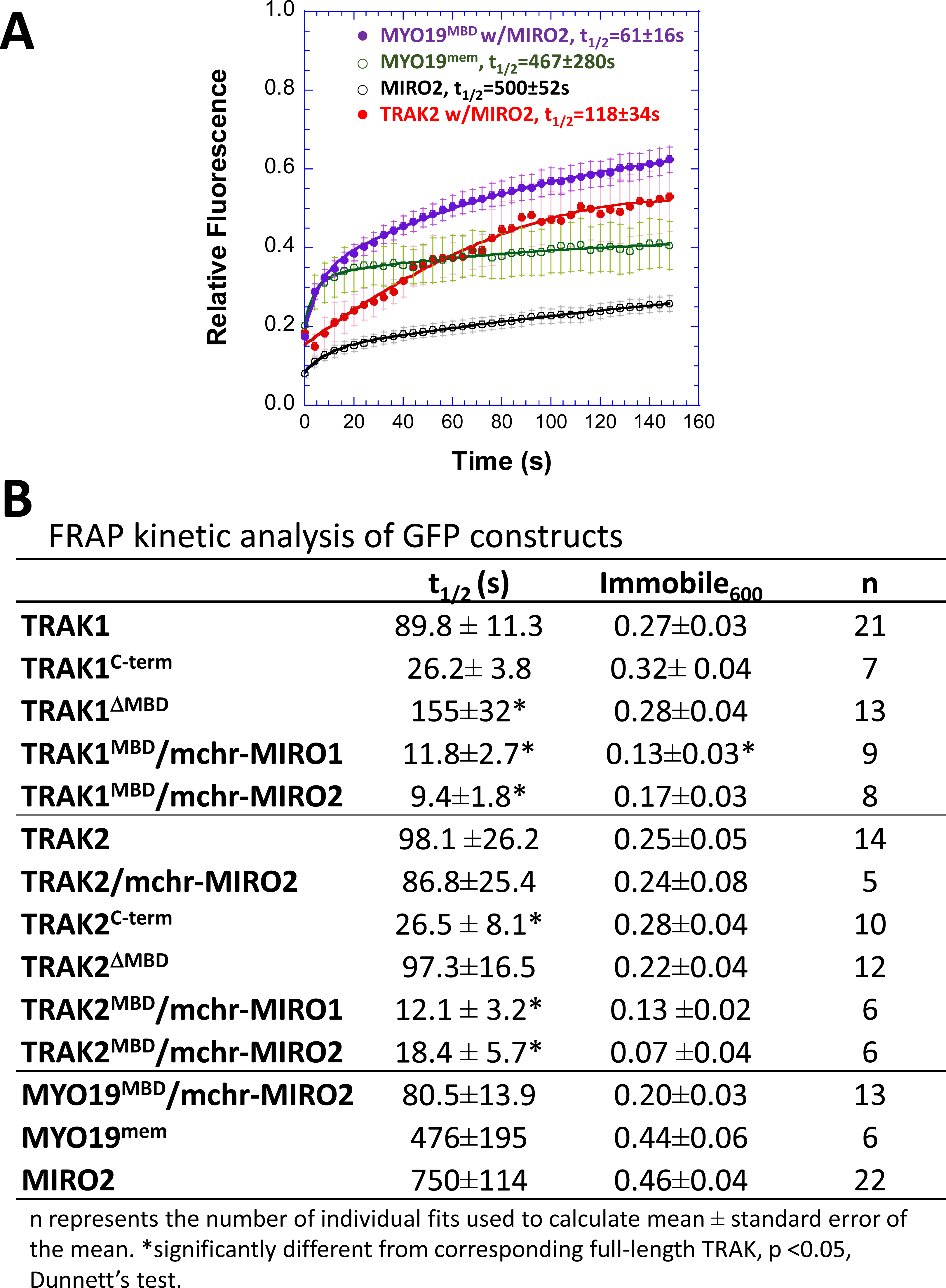
FRAP analysis reveals that GFP-MYO19 truncation construct’s exchange kinetics are slower than the kinetics for the corresponding GFP-TRAK constructs. (A) When expressed in MEF^dko^ cells, GFP-MYO19^MBD^ in combination with mchr-MIRO2 displayed slower exchange kinetics than what was observed for GFP-TRAK^MBD^ constructs coexpressed with mchr-MIRO1 or mchr-MIRO2. GFP-MYO19^mem^, which also displays MIRO-independent mitochondrial localization, had a t .20-fold longer than the corresponding GFP-TRAK^C-term^ constructs. GFP-TRAK2 coexpressed with mchr-MIRO2 displayed similar FRAP recovery kinetics to conditions where mitochondrial localization was MIRO2-independent. Points represent the mean ± standard error at each time point, and the curves are a double-exponential rise calculated from the averaged data, n>= 5 cells. (B) Rather than representing the fits calculated from the averaged data at each time point like in Figure 4 and S3A, the data in the table represent the descriptive statistics calculated from averaging the kinetic parameters from the number of fits to individual cells, represented by n for each condition.

**Figure S5:**
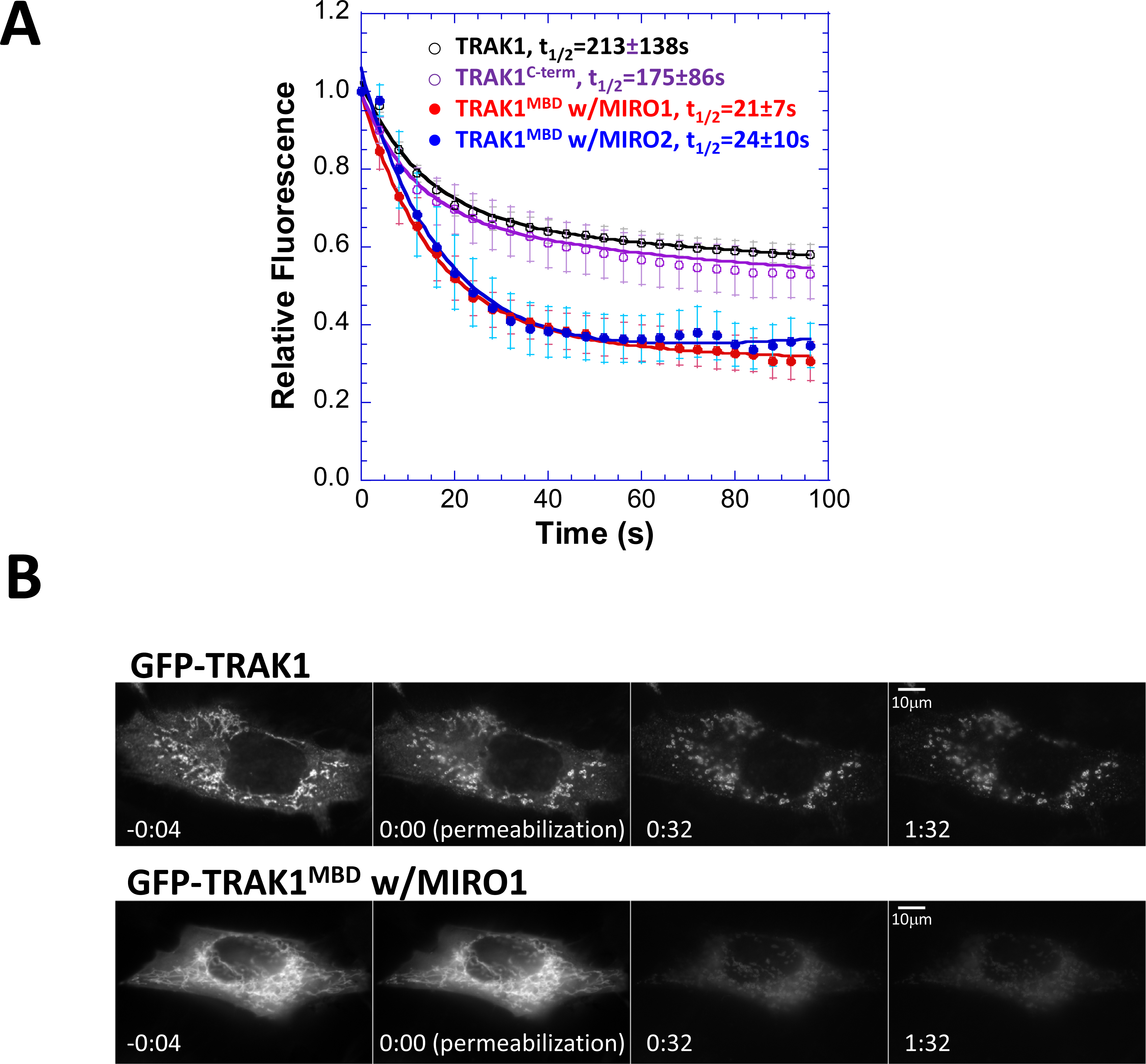
Permeabilization reveals that GFP-TRAK1 dissociates from mitochondrial pools more slowly than GFP-TRAK1^MBD^ associated with mchr-MIRO1. (A) When expressed in MEF^dko^ cells, GFP-TRAK1^MBD^ in combination with mchr-MIRO1 displayed faster fluorescence loss from mitochondrial regions than either GFP-TRAK1 or GFP-TRAK1^C-term^. Points represent the mean ± standard error at each time point, and the curves are a double-exponential decay calculated from the averaged data, n>= 9 cells. (B) Representative imaging series for PARF samples. GFP-TRAK1^MBD^ fluorescence dissipated more rapidly from MEF^dko^ cells expressing mchr-MIRO1 than GFP-TRAK1 did from MEF^dko^ cells that were not expressing either MIRO. The patterns observed in these images are representative of the patterns observed in cells expressing different combinations of constructs but with similar PARF kinetics. Time is indicated in minutes:seconds.

**Figure S6:**
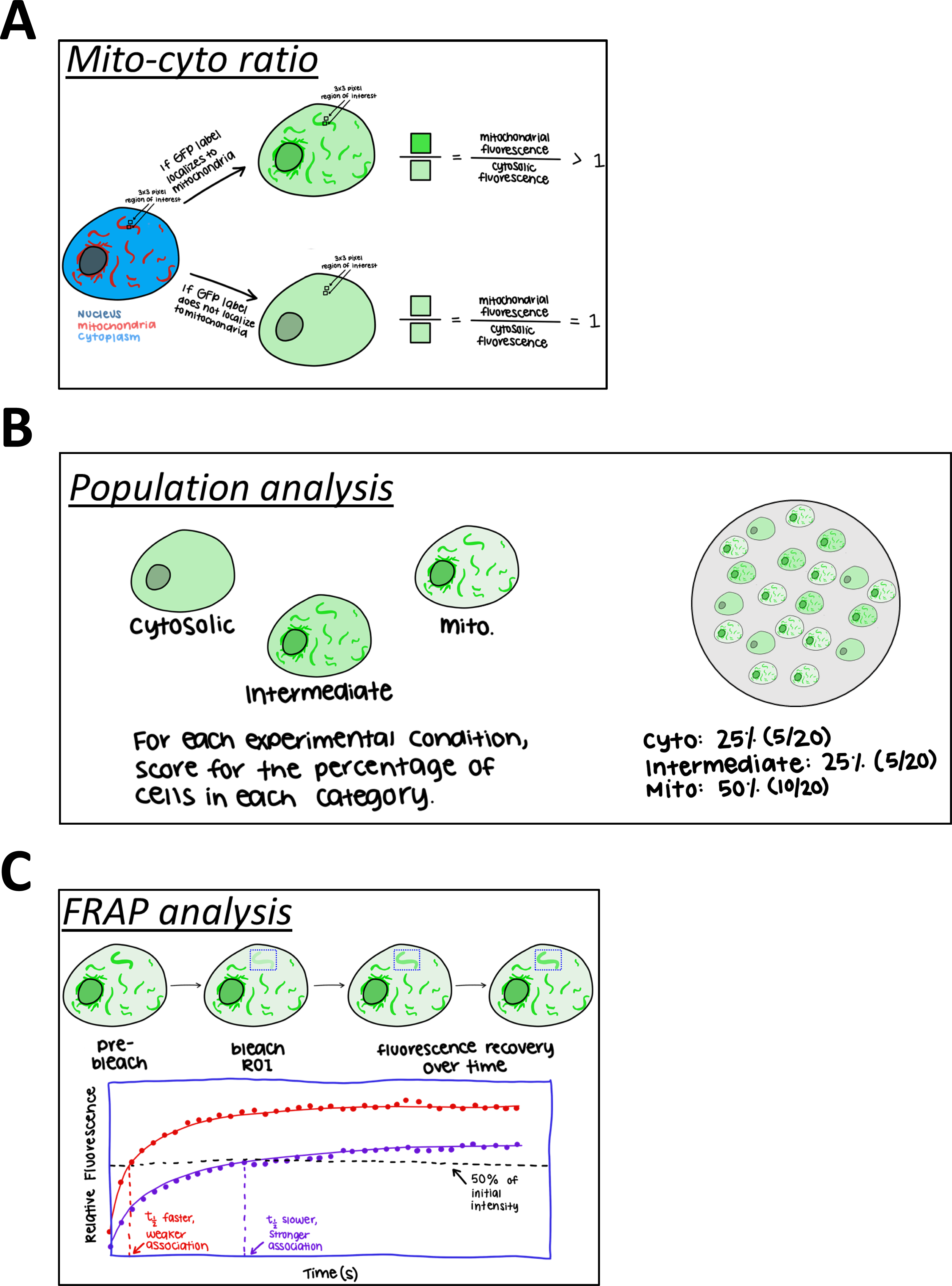
Schematics of the quantitative imaging approaches used. (A) The ratio of mitochondrial fluorescence to cytoplasmic fluorescence was calculated by dividing the mean fluorescence signal for the GFP channel in a ROI within the mitochondria by the mean fluorescence for the GFP channel in an adjacent cytoplasmic region. Five such measurements were averaged together for each cell. (B) To determine the fraction of cells in a population displaying a particular phenotype (mitochondrial, cytosolic, or intermediate), the number of cells with a particular phenotype were divided by the total number of cells counted for that experimental condition. (C) Fluorescence recovery after photobleaching analysis involved photobleaching a region of interest and then collecting a time-lapse series following the photobleaching. The brightness value of the bleached ROI was normalized to the brightness value of that same ROI in the frame prior to bleaching, and corrected for photofading due to acquisition. The resulting data were then fit to a double-exponential rise, and the time to 50% of the initial brightness (t_1/2_) and the immobile fraction were calculated from the double-exponential fit. Longer t_1/2_ were interpreted as slower exchange of bleached fluors for unbleached fluors, indicating a stronger association of that particular fluor with its associated subcellular assembly.

## Materials and Methods

### Construct generation

Human mchr-MIRO1, mchr-MIRO2 (Bocanegra et al., 2019), and GFP-MIRO2 (Birsa et al., 2014) were previously reported. Mouse GFP-TRAK constructs used in these studies were generated using previously reported GFP-TRAK plasmids (Lopez-Domenech et al., 2018) as a template, after the addition of an additional sequence of amino acids to the linker between the GFP and TRAK. Truncations were generated using PFU Ultra II PCR enzyme for megaprimer insertion PCR or deletion mutagenesis PCR, depending on the nucleotide changes required (Geiser et al., 2001), or ordered from Twist Bioscience. A schematic of the constructs used can be found in Figure S1.

### Cell culture and transfection

HeLa cells (Scherer et al., 1953) and MIRO1/2 double-knockout mouse embryonic fibroblasts (MEF^dko^) (Lopez-Domenech et al., 2018) were grown in DMEM high glucose (ThermoFisher) supplemented with 10% fetal bovine serum (Gemini Bio-products) 50 units/mL penicillin, and 50μg/mL streptomycin. Cells were maintained in a humidified incubator at 37°C and 5%CO_2_. Cells were passaged using 0.25% trypsin-EDTA. For imaging experiments, HeLa cells were grown on 22mm #1.5 glass coverslips. For live-cell experiments MEF^dko^ cells were grown on #1.5 coverslip glass-bottom 35mm dishes.

Cells were transfected with Lipofectamine 3000 (ThermoFisher) using a modified manufacturer′s protocol. For all experiments, 3μL of enhancer reagent, 1μg of GFP-tagged construct DNA were mixed with 0.5μg of mchr-tagged construct DNA and diluted in Optimem (ThermoFisher) without serum or antibiotics, in a final volume of 125μL per coverslip. In all experiments, 4μL of Lipofectamine was diluted into 125μL of Optimem without serum or antibiotics and then mixed with the DNA dilution. Complexes were allowed to form for 5 minutes at room temperature. The entirety of the DNA/reagent mix was added drop-wise to a well of a 6-well plate or one glass-bottom dish. Cells were used for experimentation 14 to 30 hours after transfection.

### Cell fixation and image acquisition

Transfected cells grown on coverslips were stained with 100nM Mitotracker DeepRed FM diluted in growth media for 10 minutes, and then washed in growth media for 10 minutes. Cells were then fixed in PBS with 4% paraformaldehyde 37°C. After permeabilization in PBS with 0.5% TritonX-100 for 5 minutes, cells were stained in PBS with 3.3 nM ALEXA plus 405-phallodin for 20 minutes to label the actin cytoskeleton. Coverslips were washed in PBS four times for five minutes and mounted on slides using PBS in 80% glycerol and 0.5% N-propyl gallate. Coverslips were sealed onto slides with Sally Hansen Hard as Nails clear nail polish.

Images were acquired using an Olympus IX-83 microscope and a PLAN APON 60x/1.42NA DIC objective. Cells were illuminated using an EXFO mixed gas light source, Sedat Quad filters (Chroma), and Sutter filter wheels & shutters. Images were acquired using a Hamamatsu ORCA-Flash 4.0 V2 sCMOS camera, and Metamorph imaging software to control the system components. In all instances, exposure times maintained constant across experimental data sets.

### Mito/Cyto Ratio calculation

The ratio of mitochondria localized fluorescence to cytosol localized fluorescence (mito/cyto ratio, MCR) was calculated by measuring the mean gray value of mitochondria and cytosol with 3×3 pixel boxes using FIJI software (Schindelin et al., 2012). The ratio was calculated for five separate regions within one cell and then averaged across the cell.

The assumption of the MCR is that GFP-TRAK fluorescence will evenly disperse throughout the entire cell unless TRAK is specifically targeted to a cellular location, such as the mitochondria. “Mitochondrial regions” of cells were identified by using the Mitotracker channel. Mitochondrial boxes were placed where mitochondria were visible. Boxes were placed for “cytosolic regions” adjacent to mitochondrial boxes in areas of low Mitotracker fluorescence where no mitochondria were present. Measurements were not obtained from the Mitotracker channel. Instead, mito and cyto boxes placed in the Mitotracker channel were applied to corresponding images from the “GFP-TRAK” channel. The fluorescence of the mitochondria relative to the fluorescence of the cytosol in the GFP-TRAK channel thus indicated the relative concentration of TRAK proteins in mitochondrial regions compared to the cytoplasm. An MCR that has a value greater than 1 indicates that mitochondria-associated GFP-TRAK fluorescence was greater than cytosolic GFP-TRAK fluorescence. An MCR equal to 1 indicates that TRAK was not localizing differently between the mitochondria and the cytoplasm (see Figure S6A for a schematic).

### Population analysis of phenotypes

Phenotypic variability exists in any population of cells or organisms. Cells were visually scored to identify the prevalence of strongly localized GFP-TRAK across the population of cells by assigning cells to one of three categories: cytosolic, mitochondrial, or intermediate localization (Bocanegra et al., 2019). Cells characterized as “cytosolic” showed no distinct GFP-TRAK fluorescence in mitochondrial regions, while cells characterized as “mitochondrial” displayed strong fluorescence in mitochondrial regions with little fluorescence in cytosolic regions. “Intermediate” cells exhibited strong fluorescence in mitochondrial regions in conjunction with some fluorescence in cytosolic regions. During population scoring, the data collector was blinded to the identity of the samples. The fraction of cells exhibiting each TRAK localization phenotype was calculated by manually scoring 60-100 cells per coverslip. The percentage of cells of each phenotype was calculated by dividing the number of cells of a specific phenotype by the total number of cells counted for the coverslip (see Figure S6B for a schematic).

### Fluorescence recovery after photobleaching (FRAP) analysis

Cells were grown on coverslip-glass-bottom dishes prior to transfection as previously described. For live-cell imaging, growth media was exchanged for imaging media (Optimem without phenol red containing 50 units/mL penicillin, and 50μg/mL streptomycin), and cells were maintained in a humidified environment at 37°C and 5% CO_2_ using a stage-top OKO-labs incubation chamber. Images of cells displaying strong mitochondria localization for the GFP-construct of interest were collected on an Olympus FV1200 laser scanning confocal microscope outfitted with a PLAN APON 60x/1.4NA objective at a frame-rate of 1 frame every 4 seconds. After the first 15 frames, regions of interest were illuminated for photobleaching at high laser power for 1 second. FIJI was used to identify regions of interest and calculate average fluorescence intensity for each time point. Relative fluorescence at each time point was measured by determining the fluorescence intensity relative to the frame prior to photobleaching (t = -4s). Data were corrected for photofading due to imaging (Applewhite et al., 2007) and averaged together. The mean gray value relative to t = -4s was plotted over time and fit to the function y = a*(1 - *e*^−bt^) + c*(1 - *e*^−dt^) +e. The half-life (t1/2) was calculated using from the exponential rise function by calculating the value of t when y = 0.5 (see Figure S6C for a schematic). Immobile fraction was calculated by determining the asymptote being approached at time = 600s. Regions of interest chosen for analysis were of sections of a reticular mitochondrial network, rather than of individual mitochondrial particles. Additionally, ROI were drawn to minimize area devoid of mitochondria whenever possible.

### Permeabilization activated reduction in fluorescence (PARF) analysis

The experimental procedure for PARF analysis was previously described (Singh et al., 2016). Briefly, cells were grown on coverslips prior to transfection as previously described. Coverslips were mounted in an open-top Rose chamber (Rose et al., 1958) in 300 μl KHM buffer (pH 7.4, 110 mM potassium acetate, 20 mM HEPES, 2 mM MgCl_2_), and maintained at ∼35°C on the microscope stage using a Nevtek air-curtain. Images of cells displaying strong mitochondria localization for the GFP-construct of interest were collected on an Olympus IX-83 microscope with a PLAN APON 60x/1.42NA DIC objective. Cells were illuminated using an EXFO mixed gas light source, Sedat Quad filters (Chroma), and Sutter filter wheels & shutters. Images were acquired using a Hamamatsu ORCA-Flash 4.0 V2 sCMOS camera at an interval of one image every four seconds. After the first 20 frames, digitonin was added (t = 0) to a final concentration of 25 μM. FIJI was used to identify regions of interest and calculate average fluorescence intensity for each time point. Fluorescence intensity relative to the frame prior to permeabilization (t = -4) was calculated for multiple experiments. The mean gray value relative to t = -4 was plotted over time and fit to a double exponential decay function: *y* = *a* + *b*\**e*^(−ct)^ + *d*\**e*^(−ft)^. The half-life (t_1/2_) was calculated as the value of t when *y* = 0.5.

### Quantification, analysis, and statistics

Metamorph (Universal Imaging) and FIJI were used for image analysis. Data points are expressed as mean ± standard error of the mean. Data were compared by Student’s t-test or Dunnett’s analysis using Kaleidagraph. Exponential fits for FRAP or PARF analysis were performed using Kaleidagraph. Parameters calculated from exponential fits in the data table are expressed as mean ± the SEM calculated across individual samples. Parameters shown on the graphs are calculated as mean ± the SEM -adjusted fit (Singh et al., 2016), by generating curve fits of the mean + SEM and mean − SEM, and reporting the error as the difference between the value calculated from the mean fit and the value calculated from the SEM-adjusted fit. All images were prepared for publication using FIJI, Metamorph, Photoshop, or some combination of these software packages.

## Notes

### Competing Interest Statement

The authors have declared no competing interest.

### Summary of Updates

Added corresponding MIRO1 experiments, added quantification of ectopic expression levels, added clearer descriptions of FRAP analysis and representative images, added PARF analysis and representative images. Supplemental images in main document.

